# Gut bacterium of *Leishmania donovani*-non-transmitting sand fly induces IL-23∼Th17 axis to protect hosts from drug-resistant visceral leishmaniasis

**DOI:** 10.64898/2026.10.04.756455

**Authors:** Suman Karmakar, Pritam Mandal, Supriya Nath, Soumyadip Mukherjee, Kamalika Roy, Mintu Karan, Sharmistha Paul, Sondipon Chakraborty, Ashif Ali Sardar, Mithun Sarkar, Pradyumna Patra, Chiranjib Pal

**Author notes:** Equal contribution (Pritam Mandal, Supriya Nath, Soumyadip Mukherjee, Kamalika Roy). Corresponding author: Chiranjib Pal, Cellular Immunology and Vector Molecular Biology Laboratory, Department of Zoology, West Bengal State University, Barasat, North 24 Parganas, Pin-700126, West Bengal, India.

## Abstract

**Background:** Visceral leishmaniasis (VL) is caused by *Leishmania donovani* and differentiates under the influence of biotope-specific gut bacteria in the *Leishmania donovani*-transmitting sand fly vector, *Phlebotomus argentipes,* which coexists with the *Leishmania donovani*-non-transmitting *Sergentomyia* species in the Indian subcontinent.

**Hypothesis:** *Sergentomyia* are primarily herpetophilic. Interestingly, we identified a specific population of anthropophilic *Sergentomyia babu babu*, a proven naturally non-competent vector for transmitting *Leishmania donovani*, with high abundance and human-bite incidences in VL-non-endemic regions, compared with VL-endemic regions of India. Thus, the possibility of egesting gut bacteria to humans from VL-non-transmitting *Sergentomyia* and inducing downstream immune cascades remains undeniable.

**Innovation:** We characterized gut bacteria from anthropophilic *Sergentomyia babu babu* in both VL-endemic and VL-non-endemic regions of India and found that a heat-inactivated formulation of *Heyndrickxia sporothermodurans* [NCBI Sequence Read Archive (SRA): PRJNA1403236], isolated from anthropophilic *Sergentomyia babu babu* from a VL-non-endemic region, prevented and cured *L. donovani* infection by inducing IL-23∼Th17. More importantly, it also prevented the initial parasite establishment by inducing phagocytosis, ROS, and TNF-α in circulatory Ly6G^−^CD11b^+^Ly6C^hi^ monocytes. To the best of our knowledge, this is the first report confirming that the gut bacteria of VL-non-transmitting sand flies are critical and need to be investigated in depth for developing therapeutics and/or prophylactics against visceral leishmaniasis (VL), and suggests that possible bite-transfer of gut bacteria from disease-non-transmitting vectors to hosts may restrict disease pathogenicity. Further clinical trials are needed to warrant (Indian patent: 202511041013, April 28, 2025).

## Introduction

Vector-borne diseases are a spectrum of infectious diseases caused by parasites, bacteria, and viruses transmitted through the bites of hematophagous insects and account for more than 700,000 annual deaths.^1^ Multiple species of *Leishmania* parasites manifest three severe clinical forms: cutaneous leishmaniasis (CL), mucocutaneous leishmaniasis (MCL), and visceral leishmaniasis (VL).^2^ In the Indian subcontinent, *Leishmania donovani* causes VL and is transmitted by the VL-transmitting sand fly, *Phlebotomus argentipes*.^2,3^ However, in reality, the VL-transmitting sand fly, *Phlebotomus argentipes,* and VL-non-transmitting sand flies, *Phlebotomus papatasi* and *Sergentomyia* species, coexist in endemic and non-endemic regions in the Indian subcontinent.^4,5^ Surprisingly, the VL-transmitting *P. argentipes* are primarily zoophilic and secondarily feed on human blood, whereas the VL-non-transmitting *P. papatasi* are primarily anthropophilic and opportunistically feed on animals.^6–10^ The other coexisting *Sergentomyia* species primarily feed on cold-blooded reptiles and amphibians, but occasionally feed on mammals, including humans.^11,12^ The VL-endemic regions of the Indian subcontinent have been reported to carry *Leishmania* infection in female *P. argentipes* at levels as low as 0.468-0.578% in Nepal^13^ and as high as 4.90-17.37% in India^14^. Interestingly, *Sergentomyia* appeared as a distinct species, even higher than *P. argentipes,* in endemic and non-endemic regions in India.^14–17^ Each bite of either infected or uninfected VL-transmitting or VL-non-transmitting sand flies injects a complex mixture of salivary proteins into the hosts’ circulation, which can initiate downstream immune cascades in hosts, either anti-parasitic or pro-parasitic.^18–21^ It was also observed that vector-specific gut microbiotas egested during bites^22^ and host-specific skin commensals^23,24^ could enhance disease severity by inducing IL-1β and neutrophil recruitment during visceral and cutaneous infection. As both specific VL-transmitting sand flies and several VL-non-transmitting sand flies coexist, an individual cannot ignore the chances of getting bitten by the latter and ingesting gut microbiota in real time before, after, or even when *P. argentipes* is inoculating *Leishmania* into the host’s circulation. Several classical reports have revealed tripartite interactions among *Leishmania*, the microbiota, and sand flies^25,26^; the influence of vector’s salivary components on disease pathogenicity^18–21^; and co-infection of *Leishmania* and the microbiota in hosts.^22–24^ However, the information that is entirely missing is the influence of gut bacteria of VL-non-transmitting sand flies on the pathogenicity of *L. donovani* infection. How the pre-inoculation or the post-inoculation of these gut bacteria influences the parasite infection, as the VL-non-transmitting sand flies are also capable of inoculating both salivary components and gut bacteria into hosts, except the infective *Leishmania*. Given the global demand for anti-leishmanial vaccine development, it is necessary to identify the probable gut bacteria egested by VL-transmitting or VL-non-transmitting sand flies.

Initially, we have been working on the abundances of VL-non-transmitting *Sergentomyia* sand fly species available in West Bengal, India. The hypotheses of this study originated from one of our critical observations that a specific species of *Sergentomyia* has preferential feeding on human blood in the VL-non-endemic district South 24 Parganas, West Bengal, India, which was reported to be endemic till 2014.^27^ Morphological identification^28,29^ and barcoding using the Cytochrome c oxidase I (COI) mitochondrial gene^30,31^ confirmed that the human-probing VL-non-transmitting sand fly is *Sergentomyia babu babu.* In contrast, the other available species, *S. shorttii* and *S. punjabensis*, feed on reptiles.^32^ Interestingly, the abundances of *S. babu babu* were found to be higher in the VL-non-endemic district, South 24 Parganas, in comparison to VL-endemic districts, Malda and Darjeeling^33^ (Fig. 1A, Supplementary Fig. 1). More importantly, the incidence of human probing appeared directly proportional to the higher abundance of *Sergentomyia babu babu* in VL-non-endemic districts (Fig. 1A, 1B).

**Fig. 1:**
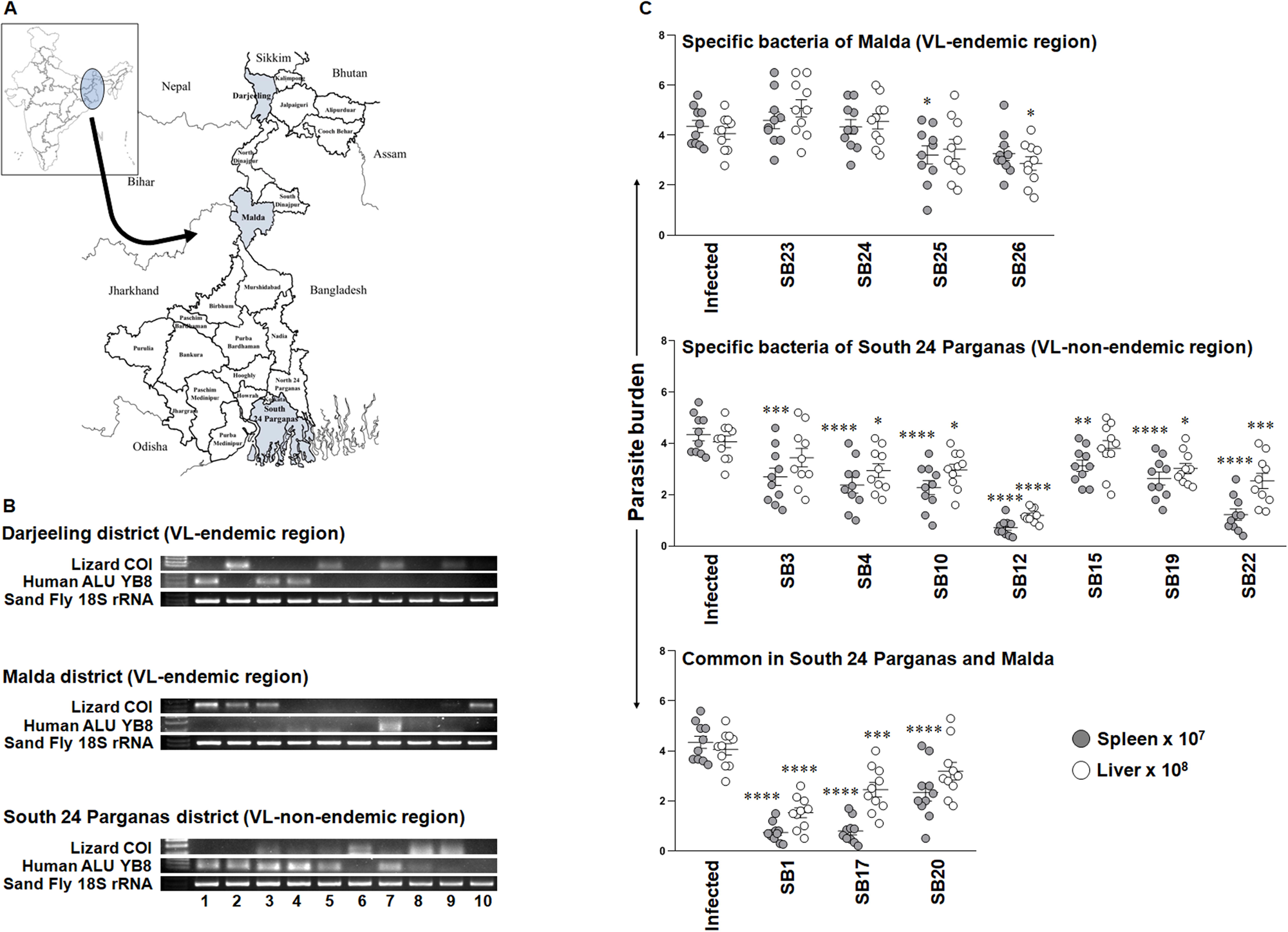
Differential anti-leishmanial effect of bacteria isolated from the gut of field-caught human-probing *S. babu babu* from VL-endemic and VL-non-endemic regions. **(A)** The wild-caught sand flies were sampled from the regions of three different VL-endemic and VL-non-endemic districts, as indicated on the map of West Bengal, India. **(B)** The blood meal of individual sand flies (n=10 in each district) was identified using both lizard (*Hemidactylus frenatus*)-specific Cytochrome Oxidase I and the human AluYB8 gene by agarose gel electrophoresis. The abundance of human-fed *Sergentomyia babu babu* was found to be high in VL-non-endemic district and low in VL-endemic districts. **(C)** Among all VL-non-endemic and VL-endemic districts’ bacterial isolates, SB1, SB12, SB17, and SB22 showed anti-leishmanial activity against antimony-resistant *L. donovani* at one month post-infection *in vivo*, as determined by Giemsa-stained micrographs and the Stauber formula. *Heyndrickxia sporothermodurans* (SB12) was found to be the most effective therapeutic lead against antimony-resistant *L. donovani, in vivo*, isolated from anthropophilic *S. babu babu* in a VL-non-endemic district. The statistical measure (mean±SEM) was used to interpret all experimental data from two independent experiments comprising of 5 biological replicates each, followed by One-way ANOVA with Dunn’s post hoc analysis to calculate the p-value. Each dot in the graph depicts the value derived from an individual experimental animal, *p<0.03, **p<0.006, ***p<0.0006, ****p<0.0001, vs PBS-treated infected control. SB1: *Bacillus licheniformis*; SB12: *Heyndrickxia sporothermodurans;* SB17: *Bacillus subtilis;* SB22: *Paenibacillus dendritiformis*. Source data file has been provided with the Source data. See also Supplementary Fig. 1, 2 and Supplementary Table 1-3.

As evidenced, the VL-transmitting sand fly, *Phlebotomus argentipes,* and VL-non-transmitting sand flies, *Phlebotomus papatasi* and *Sergentomyia* species, coexist in VL-endemic and non-endemic regions in the Indian subcontinent.^4,5^

And whereas, we identified a population of unconventional anthropophilic *Sergentomyia babu babu* with high abundance in VL-non-endemic districts of West Bengal, India.

And whereas, the bacteria, either vector-specific egested during bites^22^ or host-specific skin commensals^23,24^, could influence the severity of leishmaniasis.

We raised the question of whether the gut bacteria of human-probing *Sergentomyia babu babu* found in VL-non-endemic regions could inhibit *L. donovani* infection, in view of vaccine development against drug-resistant infection.

## Results

### Identification of wild-caught sand flies

A total of seven thousand six hundred and ninety-five wild-caught sand fly specimens were collected from three geographically separated VL-endemic and non-endemic regions of Malda, South 24 Parganas, and Darjeeling. Two genera (*Sergentomyia* and *Phlebotomus*) and four species (*Sergentomyia babu babu, Sergentomyia shorttii, Sergentomyia punjabensis,* and *Phlebotomus argentipes*) were identified morphologically based on the number of teeth recorded 25 to 30, 10 to 14, and 30, respectively, in the cibarium of *Sergentomyia* females according to Lewis’s identification key^28,29,34^ under the stereo-zoom microscope (Stemi 305, Zeiss) and inverted phase contrast microscope (Leica DMi8) (Supplementary Fig. 2A). To ensure molecular-level identification, all specimens were barcoded using the mitochondrial COI gene^30,31^, and the amplified COI sequences were submitted to the NCBI GenBank database (Table 1). The neighbour-joining tree constructed using the COI gene shows that *Sergentomyia babu babu* and *Sergentomyia shorttii* are clustered together, with strong bootstrap support, whereas *Sergentomyia punjabensis* and *Phlebotomus argentipes* represent two separate clades as the most evolutionarily divergent species^35^ (Supplementary Fig. 2B).

### Blood meal preference of wild-caught *Sergentomyia babu babu*

As the question was about the unorthodox anthropophilic or natural herpetophilic nature of *Sergentomyia babu babu*, the bloodmeal was analyzed based on the lizard (*Hemidactylus frenatus*)-specific COI gene^32,36^ and the human AluYB8 element PCR.^37^ Multiple blood-fed wild-caught sand flies were analyzed, and the percentage of human-probing *Sergentomyia babu babu* was observed inversely proportional with the availability of VL patients, i.e., it appeared high in non-endemic South 24 Parganas and low in endemic districts, Malda, and Darjeeling (Fig. 1B).

### Diversity of culturable gut bacteria from wild-caught *Sergentomyia babu babu*

As we focused on the isolation and identification of gut bacteria and on exploring the possibility of developing a vaccine against *Leishmania* infection, we selected two epidemiological conditions. South 24 Parganas appeared to be endemic until 2014; however, very few cases were reported thereafter, and Malda district, which remains endemic, reported the highest number of cases in West Bengal, India.^33^ Interestingly, the culturable gut bacteria of South 24 Parganas and Malda differ in many cases, as identified by 16S rRNA-specific Sanger sequencing, and are mentioned in Table 2.^38^ Our next question was about the influence of this diverse bacterial population on the drug-resistant *L. donovani* infection, if any.

### Differential effect of culturable gut bacteria isolated from *S. babu babu* of VL-endemic and VL-non-endemic regions

Initially, we attempted to identify the most effective bacteria by comparing gut isolates of S. babu babu from the VL-non-endemic region with those from the VL-endemic region (Fig. 1C). Four bacteria provided substantial anti-leishmanial effects against drug-resistant L. donovani in vivo in experimental animals. Two (SB1, SB17) were common for VL-non-endemic and VL-endemic regions; however, SB12, the bacterial isolate from the VL-non-endemic region, was found to be the most promising one [SB1: *Bacillus licheniformis*; SB12: *Heyndrickxia sporothermodurans;* SB17: *Bacillus subtilis;* SB22: *Paenibacillus dendritiformis* (Fig. 1C)]. The splenic parasite load was reduced by 83±1.6% (p<0.0001) by *B. licheniformis*, 83±2.3% (p<0.0001) by *H. sporothermodurans*, 81.7±1.9% (p<0.0001) by *B. subtilis*, and by 71.8±6.2% (p<0.0001) by *P. dendritiformis* in comparison to the infected control. *H. sporothermodurans* (SB12) was found to be the most promising therapeutic lead as it reduced the drug-resistant *L. donovani* infection both in the spleen and liver by 83±2.3% (p<0.0001) and 70.6±2.1% (p<0.0001), respectively (Fig. 1C, Supplementary Tables 2 and 3).

We confirmed this hypothesis by precise, real-time quantitative PCR using parasite-specific *Ld*-18srRNA, normalized to equalized murine GAPDH.^39,40^ Interestingly, the therapeutic supremacy of heat-inactivated *H. sporothermodurans* has been confirmed against drug-resistant *L. donovani* infection both in the spleen and bone marrow by 6.26±0.5 fold (p<0.0001) and 2.8±0.3 fold (p<0.0001) reduction, respectively, in the mRNA expression of *Ld*-18s rRNA (Fig. 2A, Supplementary Table 4).

**Fig. 2:**
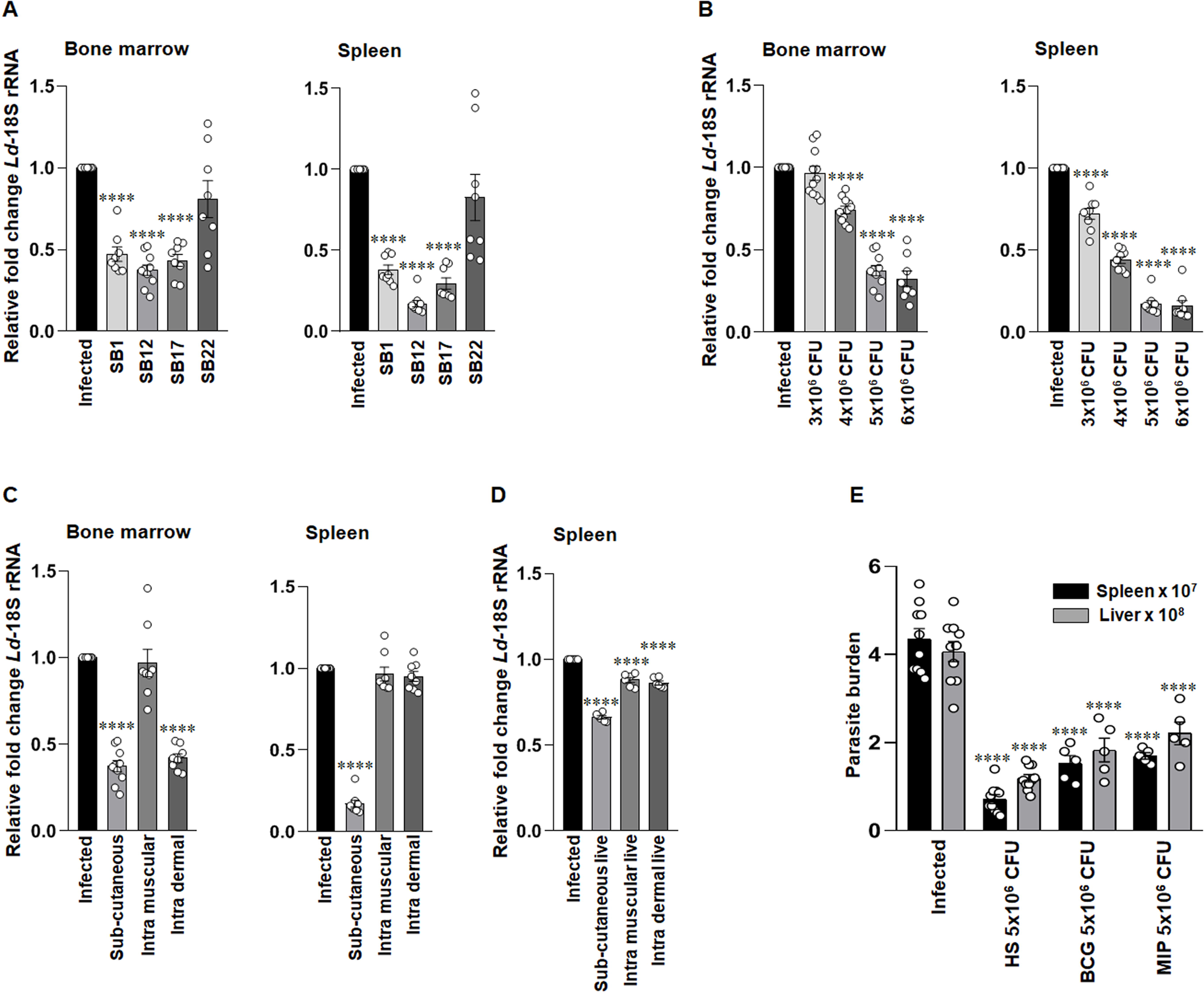
The dose selection, effective route of administration, and comparative therapeutic efficiency of heat-inactivated *H. sporothermodurans*, *in vivo*. **(A)** *H. sporothermodurans* (SB12) therapeutic dose (5×10^6^ CFU/animal) effectively reduced the parasite load in both bone marrow and spleen of treated animals, as determined by the quantification of the relative fold changes of *Ld*-18S rRNA in real-time PCR, ****p<0.0001, vs PBS-treated infected control. (B) A dose kinetics study was performed to deduce the lowest effective dose, and 5×10^6^ CFU/animal dose of heat-inactivated *H. sporothermodurans* was most effective in reducing the amastigote load in bone marrow and spleen, as determined by the quantification of the relative fold changes of *Ld*-18S rRNA in real-time PCR, ****p<0.0001 vs PBS-treated infected control. **(C, D)** Subcutaneous route of administration was found to be the most efficient route to restrict the antimony-resistant *L. donovani* (NS2) proliferation in both bone marrow and spleen of experimental animals treated with both live and heat-inactivated formulation of *H. sporothermodurans* given in subcutaneous, intradermal, and intramuscular routes after one-month post infection, as determined by the quantification of the relative fold changes of *Ld*-18S rRNA in real-time PCR,****p<0.0001 vs PBS-treated infected control **(E)** The heat-inactivated *H. sporothermodurans* showed the best therapeutic efficacy among heat-inactivated *H. sporothermodurans* (HS)*, Mycobacterium indicus pranii* (MIP), and *Bacillus*-Calmette-Guérin (BCG) in reducing the splenic and hepatic parasite load of antimony-resistant *L. donovani*, estimated by Giemsa-stained micrographs and Stauber formula, (****p<0.0001 vs infected control). Each dot in the graph depicts the value derived from an individual experimental animal. The statistical measure (mean±SEM) of minimum of 5 biological replicates combined from two independent experiments was used to interpret all experimental data, followed by One-way ANOVA with Dunn’s post hoc analysis to calculate the p-value. Source data file has been provided with the Source data. See also Supplementary Fig. 3 and Supplementary Table 4-10.

In contrast, either the abundances of human-probing *S. babu babu* (Supplementary Fig. 1) or the bite incidences (Fig. 1B) were found to be much lower in the VL-endemic region. This fact strongly established that gut bacteria from *S. babu babu,* if delivered to the host, cannot negatively affect parasite establishment. This depends on the abundance and incidence of bite exposure to hosts, which were higher in VL-non-endemic regions than in VL-endemic regions (Fig. 1B, Supplementary Tables 1 and 3). Based on this fact, we focused on developing vaccine, therapeutic, and/or prophylactic leads, specifically targeting drug-resistant *Leishmania* infections, as there is currently no licensed vaccine for human leishmaniasis.^41^

### Therapeutic dose determination and effective route of administration of *Heyndrickxia sporothermodurans*

Moving to the next step, the therapeutic dose determination of SB12, among the lower and higher doses of 3×10^6^ CFU/animal, 4×10^6^ CFU/animal, 5×10^6^ CFU/animal, and 6×10^6^ CFU/animal, 5×10^6^ CFU provided the best effect both in the spleen, liver and bone marrow and almost equivalent to 6×10^6^ CFU, and hence we rejected the latter dose to minimize any possible risk of toxicity (Fig. 2B, Supplementary Fig.3A, Supplementary Tables 5, 6).

One step further, the most essential criterion of vaccine development, either therapeutic or prophylactic, is the route of administration. Subcutaneous administration was the most effective route compared with intramuscular and intradermal administration and was effective in the spleen, liver, and bone marrow (Fig. 2C, Supplementary Fig. 3B, and Supplementary Tables 7, 8). The other point was that the heat-inactivated subcutaneous administration was far more effective than the live subcutaneous formulation (Fig. 2D, Supplementary Table 9).

Another interesting point was that heat-inactivated *H. sporothermodurans* was the most effective against drug-resistant *L. donovani* infection among the experimental therapeutic adjuvants tested, compared with Bacillus Calmette-Guérin (BCG)42,43 and *Mycobacterium indicus pranii*^44,45^ reported earlier (Fig. 2E, Supplementary Table 10).

The next question was whether the most efficient therapeutic dose would also be effective when administered on a prophylactic schedule.

### The prophylactic regimen of *Heyndrickxia sporothermodurans* prevented the viscerotropic parasite infection, *in vivo*

Importantly, the prophylactic regimen of heat-inactivated *H. sporothermodurans* prevented the drug-resistant *L. donovani* infection in both the spleen and liver. The reduction of parasite burden in the spleen and the liver was found optimistic, by 73.8±1.03% (p<0.0001) and 51±2.3% (p<0.0001) in the spleen and liver, respectively, in comparison to PBS-treated infected control (Fig. 3A, Supplementary Table 11), with further confirmation at mRNA level by a inhibition of parasite replication by 3.8±1.3 fold (p<0.0004 vs infected control) in the spleen, which confirmed that heat-inactivated *H. sporothermodurans* formulation could be used also as a prophylactic vaccine (Fig. 3B, Supplementary Table 12).

**Fig. 3:**
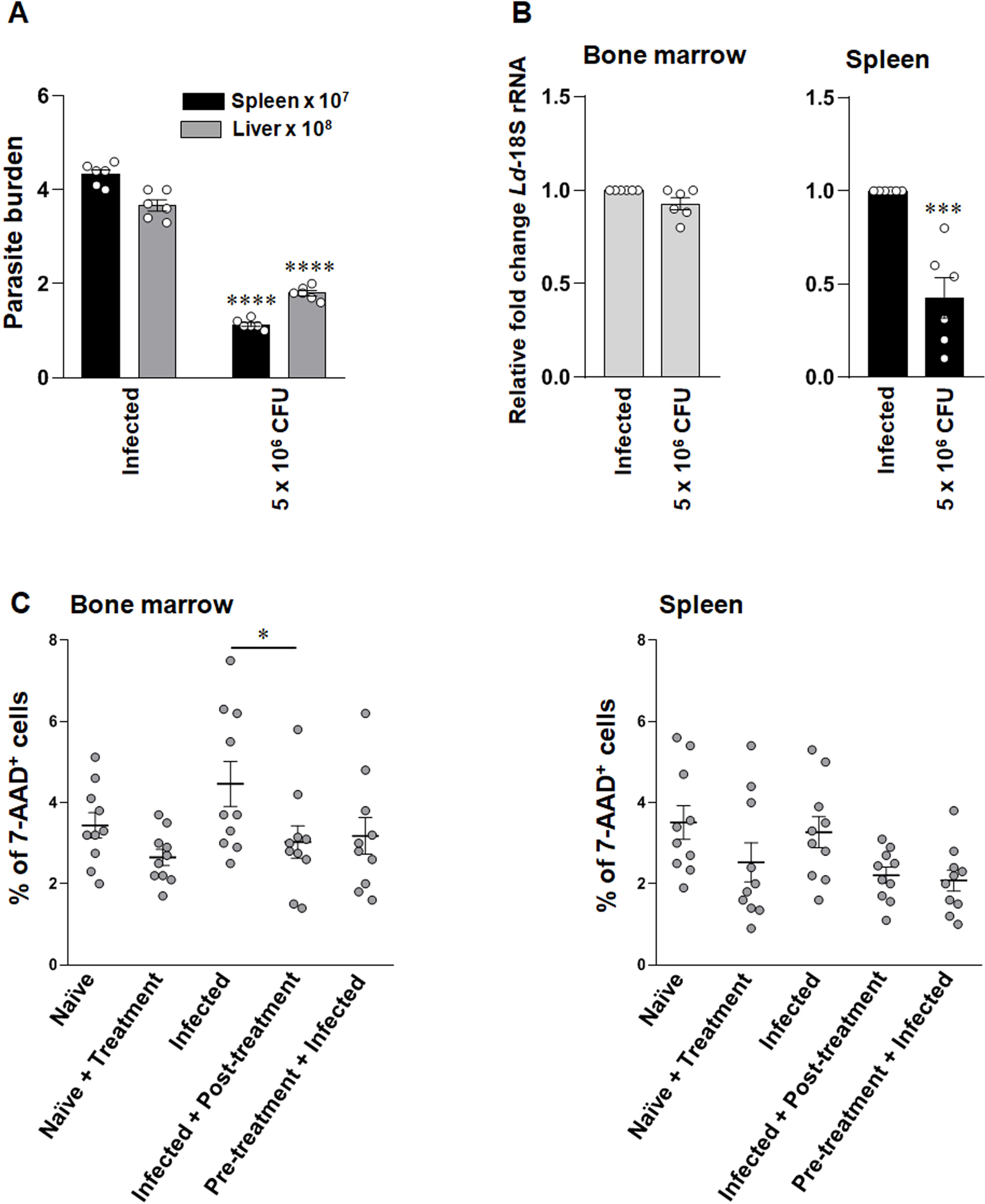
The efficacy of heat-inactivated *H. sporothermodurans* in the prevention of drug-resistant *L. donovani* and toxicity assay, *in vivo*. **(A)** The heat-inactivated *H. sporothermodurans* prophylactic treatment effectively reduced parasite proliferation in both the spleen and liver, as determined by Giemsa-stained micrographs and the Stauber formula (****p<0.0001 vs PBS-treated infected control). (B) The relative fold changes of *Ld*-18S rRNA were effectively reduced in the spleen by the prophylactic doses of 5×10^6^ CFU/ animal of heat-inactivated *H. sporothermodurans,* as determined by qPCR, ***p<0.0004 vs PBS-treated infected control. (C) The therapy and prophylactic dose (5×10^6^ CFU/ animal) of heat-inactivated *H. sporothermodurans* showed no cytotoxicity on bone marrow and splenic cells, as estimated by a cell viability assay using 7-AAD in a flow cytometer (BD FACSVerse™), *p<0.02 vs PBS-treated infected control. The statistical analyses of all the experimental groups were depicted in bar plots and scatter plots using GraphPad Prism software (v 8.0). Each dot in the graph depicts the value derived from an individual mouse. The statistical measure (mean±SEM) of minimum of 5 biological replicates combined from two independent experiments was used to interpret all experimental data, followed by Student T test (two-tailed) and One-way ANOVA with Tukey’s post hoc analysis to calculate the p-value. See also Supplementary Table 11-13.

It is worth mentioning that both the therapeutic and prophylactic strategies with this formulation remained non-toxic, as evidenced by the flow cytometric estimation of non-viable cells by 7-AAD incorporation assay in both bone marrow and spleen cells taken from the naïve and only *Leishmania*-infected animals in comparison to either heat-inactivated *H. sporothermodurans* pre-treated or post-treated animals (Fig. 3C, Supplementary Table 13)

### The cytokine cross-talk: IL-23 and IL-1β, but not IL-12, were found to be critical for anti-*Leishmania* immunity

IL-12 is believed to be the key cytokine that induces anti-parasitic IFN-γ^+^T cell responses, and high levels promote the differentiation of FoxP3^−^ Tregs to limit inflammation and tissue damage during *Leishmania* infection.^46^ Surprisingly, we observed a reduction in expression of both subunits of IL-12, IL-12p35 by 1.63±0.1 fold (p<0.0001, vs naïve) and 1.31±0.1 fold (vs infected), and also of IL-12p40 by 1.3±0.1 fold (p<0.006, vs naïve) during the therapeutic strategy (Fig. 4, Supplementary Table 14), leaving a genuine question that if not IL-12, then who is initiating the protective immunity? Interestingly, IL-23p19 was induced by 2.3±0.2 fold (p<0.0001 vs infected) and by 1.74±0.1 fold (p<0.0006 vs naïve). IL-23, the other member of the IL-12 cytokine family, can also favor pro-inflammation by inducing Th17 cells.^47^ The dominance of IL-23 over IL-12 was confirmed by the most significant induction of IL-17A (by 5.96±0.26 fold, p<0.0001, vs infected and 3±0.1 fold, p<0.0001, vs naïve). In contrast, the minimum amount of IFN-γ expression was also detected (induction by 1.35±0.14fold, p<0.04, vs infected and 1.5±0.2fold, p<0.02, vs naïve) during heat-inactivated *H. sporothermodurans* therapy (Fig. 4, Supplementary Table 14). IL-23-induced Th17 generation has been reported to be instrumental in the defense against *Leishmania*^48^, *Plasmodium*^49^, and *Trypanosoma*^50^ infections. The IL-23 and IL-1β cytokine milieu can also promote Th17 differentiation ^51–53^. We observed a steady induction of IL-1β (by 5.7±0.33fold, p<0.0001 vs infected; and 5.5±0.3fold, p<0.0001 vs naïve), which also correlated with IL-17A induction (Fig. 4, Supplementary Table 14).

**Fig. 4:**
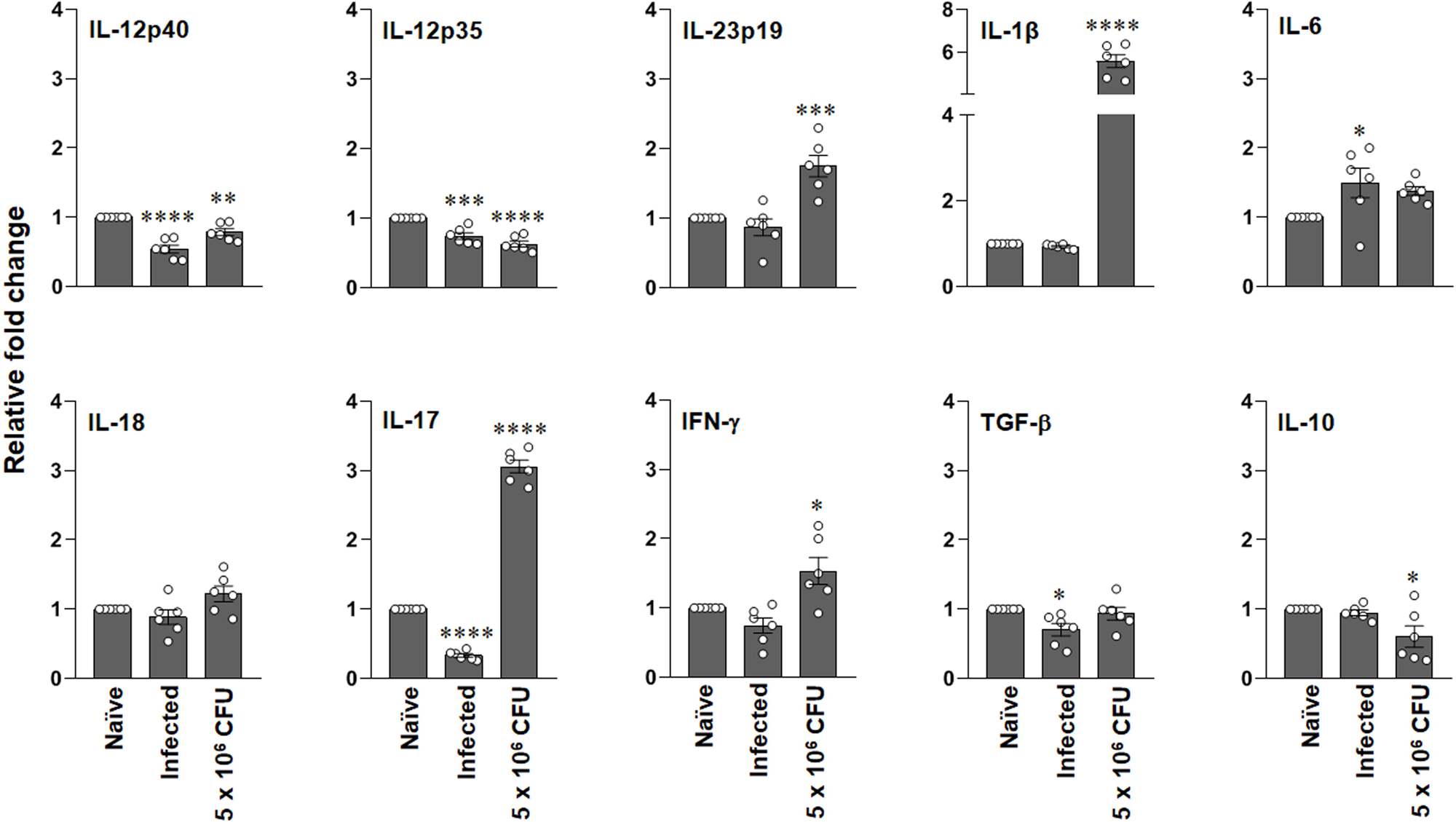
Heat-inactivated *H. sporothermodurans* induced the IL-23∼Th17 axis to protect hosts from drug-resistant *L. donovani* infection. Heat-inactivated *H. sporothermodurans* therapy induced expression of IL-1β and IL-23 and reduced the expression of both IL-12 subunits (IL-12p40 and IL-12p35) in post-treated animals compared with the other experimental groups (naïve and *Leishmania*-infected), as determined at the transcription level by real-time PCR. The significant induction of IL-17 at the mRNA level indicated that the IL-23-mediated IL-17 pathway is instrumental in combating antimony-resistant *L. donovani* infection. The proinflammatory cytokines IL-18 and IFN-γ were also induced by heat-inactivated *H. sporothermodurans* therapy, as determined by real-time PCR. Each dot in the graph depicts the value derived from an individual mouse. The statistical measure (mean±SEM) was used to interpret all experimental data from two independent sets of experiments comprising groups of 3 animals each, followed by One-way ANOVA with Dunn’s post hoc analysis to calculate the p-value. *p<0.02, **p<0.006, ***p<0.0006, ****p<0.0001 vs naïve. See also Supplementary Fig. 4, Supplementary Table 14 and 15.

Flow cytometry analysis confirmed the differential mRNA expression and showed that functional dimmers of IL-17 and IFN-γ were present. Significantly, the therapy induced the expansion of both CD8^+^IL-17^+^ T cells by 18±1.3fold, p<0.0001, and CD4^+^IL-17^+^ T cells by 6.45±1fold, p<0.0008 (Fig. 5, Supplementary Tables 16, 17). In contrast, a much smaller IFN-γ^+^ population was induced in both CD8^+^ (3.83±0.15fold, p<0.0001) and CD4^+^(2±0.14fold, p<0.0008) T cells (Fig. 5, Supplementary Tables 16, 17) compared to infected controls. Interestingly, the cytokine profiles of both therapeutic and prophylactic measures were shown to be compatible.

**Fig. 5:**
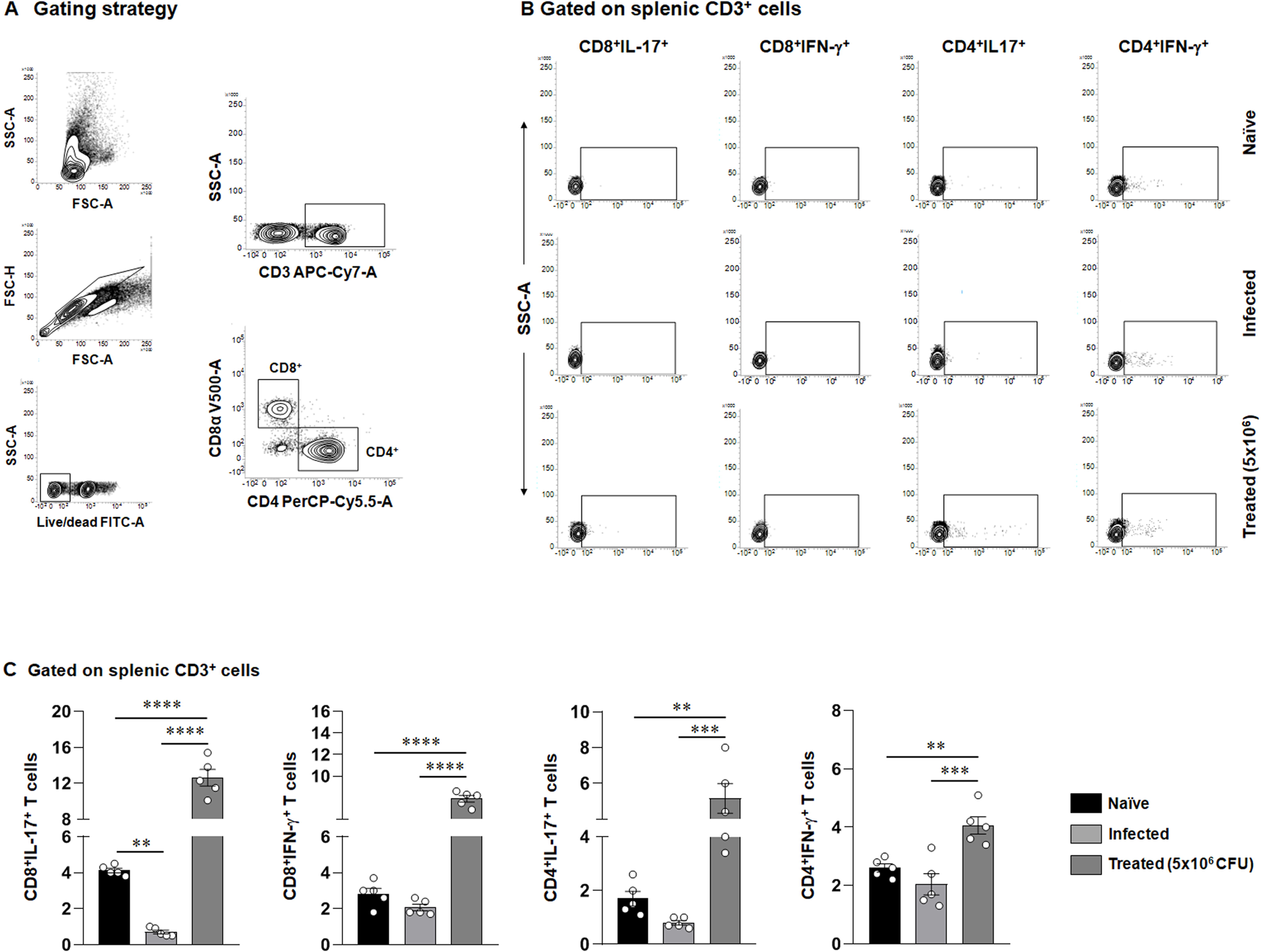
The heat-inactivated *H. sporothermodurans* therapy induced the expansion of CD8^+^IL-17^+^ T cells, CD4^+^IL-17^+^ T cells, and CD8^+^ IFN-γ^+^ T cells, CD4^+^ IFN-γ^+^ T cells in the spleen, *in vivo.* **(A)** Representative flow cytometric contour plots of CD8^+^ and CD4^+^ T cells were analyzed by gating on viable CD3^+^ cells. (B) The heat-inactivated *H. sporothermodurans* therapy induced the expansion of CD8^+^IL-17^+^, CD4^+^IL-17^+^, CD8^+^IFN-γ^+^, and CD4^+^IFN-γ^+^ T cells in post-treated experimental groups in comparison to the other experimental groups (naïve and *Leishmania*-infected), as represented by flow cytometric contour plots with respect to the isotype controls (Supplementary Fig. 6). (C) Statistical analyses were performed to plot the cumulative data for expansion of CD8^+^IL-17^+^, CD4^+^IL-17^+^, CD8^+^IFN-γ^+^, and CD4^+^IFN-γ^+^ T cells, which were represented in bar plots for all experimental groups. Interestingly, the therapy induced greater proliferation of IL-17^+^ T cells than of IFN-γ^+^ T cells in the spleen (**p<0.008, ***p<0.0008, ****p<0.0001). The statistical measure (mean±SEM) of minimum of 5 biological replicates combined from two independent experiments was used to interpret all experimental data, followed by One-way ANOVA with Tukey’s post hoc analysis to calculate the p-value. Each dot in the graph depicts the value derived from an individual experimental animal. Flow cytometric acquisition was performed using a BD FACSVerseTM, and data matrices were analyzed using BD FACSuite software. See also Supplementary Fig. 5,6 and Supplementary Table 16,17.

The supremacy of heat-inactivated subcutaneous administration over live subcutaneous formulation (Fig. 2D) can be easily explained by the higher induction of IL-23, IL-1β, and, consequently, IL-17 by the former (Supplementary Fig. 4, Supplementary Table 15).

Because both therapeutic and prophylactic approaches were effective against drug-resistant Leishmania, we were curious whether heat-inactivated *H. sporothermodurans* could block the initial entry and establishment of Leishmania in hosts.

### The prophylactic strategy activated the CD11c^−^CD11b^+^Ly6G^−^Ly6C^hi^ monocytes, resulting in disruption of the initial establishment of *Leishmania*

In real time, hosts are either pre-exposed or post-exposed to VL-non-transmitting sand fly *S. babu babu*, and may be opportunistically inoculated with *H. sporothermodurans* in circulation. The functional state of circulating monocytes is critical for initial establishment in hosts, as neutrophils transfer the parasites.^22^ Human classical monocytes (CD14^++^CD16^−^) were reported to inhibit the initial establishment of infective promastigotes by generating reactive oxygen species (ROS)^54^, and the intermediate (CD14^++^CD16^+^) monocytes were the main producers of TNF-α^55^ during CL; in contrast, monocytes of VL patients were found to have impaired phagocytic activity, intracellular ROS production, and enhanced antiinflammatory signatures.^56^ Immune activation of circulatory monocytes towards a proinflammatory signature may be a suitable approach to block the initial entry of *Leishmania*.

Interestingly, we observed the murine counterpart of inflammatory and phagocytic monocytes^57–59^, CD11c^−^CD11b^+^Ly6G^−^Ly6C^hi^ cells, to exhibit efficient phagocytic activity by 1.27±0.03 fold (p<0.005, vs infected), and interestingly, high induction of ROS by 1.8±0.05-fold (p<0.005, vs naïve) in comparison to both infected, 1.12±0.1-fold (vs naïve) and only pre-treated by 1.21±0.1-fold (vs naïve) during the prophylactic administration of heat-inactivated *H*. *sporothermodurans* and infection of the experimental animals with *Leishmania, iv,* thereafter (Fig. 6, Supplementary Tables 18, 19).

**Fig. 6:**
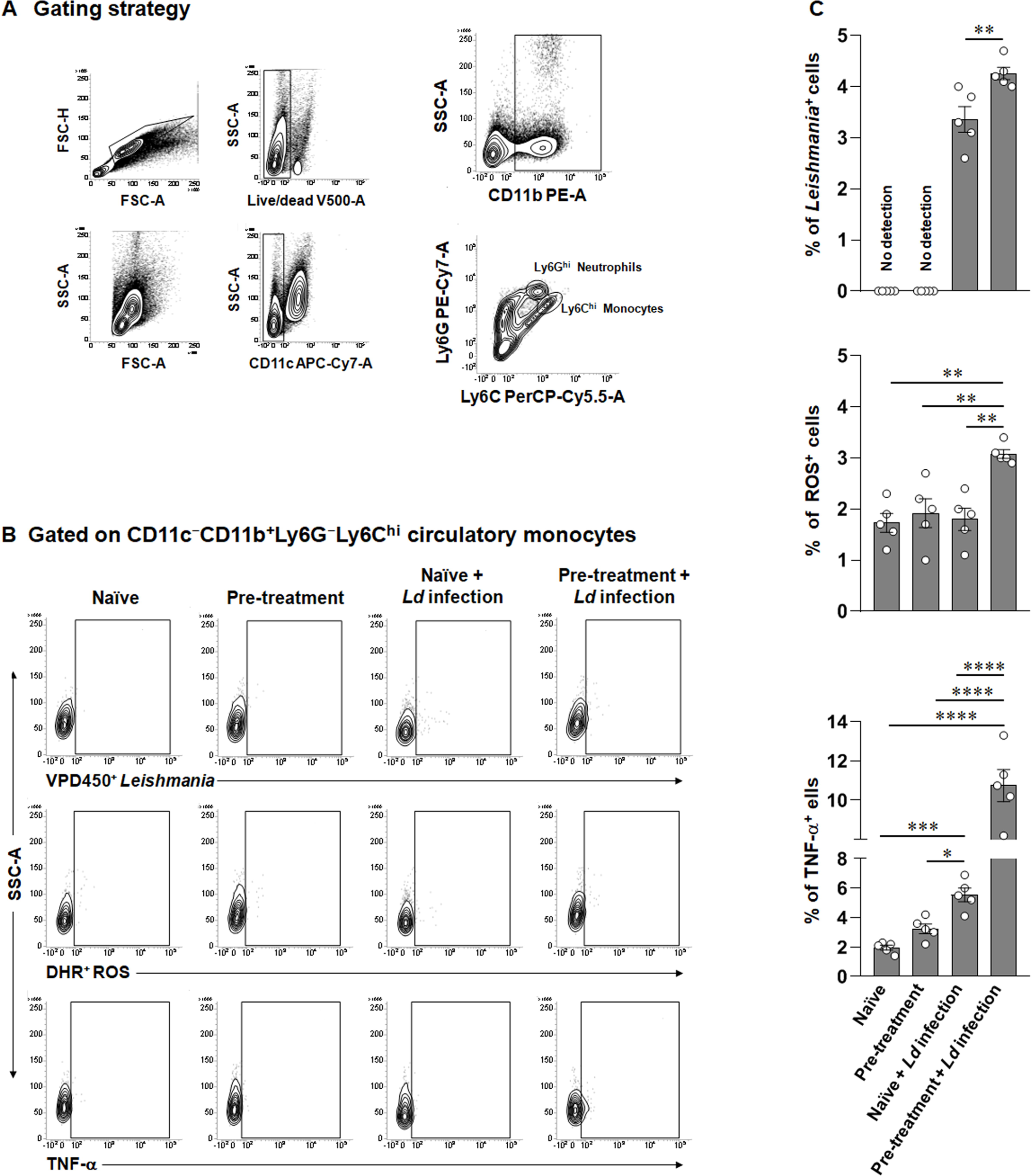
The prevention of the disease by heat-inactivated *H. sporothermodurans* involved the phagocytic uptake, oxidative burst, and release of TNF-α by circulatory CD11c^−^CD11b^+^Ly6G^−^Ly6C^hi^ monocytes, *in vivo*. **(A)** The representative flow cytometric contour plots of Ly6G^−^Ly6C^hi^ circulatory monocytes were analyzed by gating on viable CD11c^−^CD11b^+^ cells, followed by gating based on FSC-H/FSC-A contour plot on all events. **(B)** The heat-inactivated *H. sporothermodurans* pre-treatment induced the phagocytic engulfment of VPD450^+^*Leishmania* promastigotes by CD11c^−^CD11b^+^Ly6G^−^Ly6C^hi^ circulatory monocytes, ROS as determined by DHR123 oxidation, and the production of TNF-α from CD11c^−^CD11b^+^Ly6G^−^Ly6C^hi^ circulatory monocytes, as represented by flow cytometric contour plots of naïve, naïve+pre-treatment, *Leishmania*-infected, and pre-treated + *Leishmania-*infected experimental groups with respect to the unstained controls and isotype controls (Supplementary Fig. 8). **(C)** Statistical analyses were performed to plot the cumulative data of phagocytic engulfment of VPD450^+^*Leishmania* promastigotes by CD11c^−^CD11b^+^Ly6G^−^Ly6C^hi^ circulatory monocytes, high amount of ROS as determined by DHR123 oxidation, and TNF-α expression by CD11c^−^CD11b^+^Ly6G^−^Ly6C^hi^ circulatory monocytes in bar plots of all the groups of experimental animals. *p<0.02, **p<0.005, ***p<0.0007, ****p<0.0001. The statistical measure (mean±SEM) of minimum of 5 biological replicates combined from two independent experiments was used to interpret all experimental data, followed by One-way ANOVA with Tukey’s post hoc analysis to calculate the p-value. Each dot in the graph depicts the value derived from an individual experimental animal. Flow cytometric acquisition was performed using a BD FACSVerseTM, and data matrices were analyzed using BD FACSuite software. See also Supplementary Fig. 7, 8 and Supplementary Table 18, 19.

Ly6C^hi^ monocytes have been reported to be recruited during infection with broad-spectrum pathogens, including *Listeria monocytogenes*, *Toxoplasma gondii*, and *Leishmania major*^57^. Ly6C^hi^TNF-α^+^ monocytes were also found to play a key role in controlling *Toxoplasma* and *Listeria* infection^57,60,61^. We observed a high expansion of circulatory CD11c^−^CD11b^+^Ly6G^−^Ly6C^hi^ TNF-α^+^ monocytes by 5.44±0.4-fold (p<0.0001, vs naïve) in comparison to 1.65±0.2-fold (pre-treated, vs naïve) and by 2.8±0.2-fold (p<0.0007; infected, vs naïve) during the prophylactic administration of *H*. *sporothermodurans* and infection of the experimental animals with *Leishmania, iv,* thereafter (Fig. 6, Supplementary Table 19).

## Discussion

Vector salivary components have been well studied for their influence on the pathogenicity of vector-borne infectious diseases in hosts^3,18,20,62^; however, the influence of vector commensals on hosts has not been extensively studied and warrants greater attention in preventive or therapeutic interventions. Although egested gut bacteria co-deposited with parasites by VL-transmitting sand flies have been shown to enhance disease pathogenicity^22^, what is entirely missing is the influence of gut bacteria of VL-non-transmitting sand flies, as the latter are egested solely into hosts, not co-deposited with parasites^63^.

The concept of developing preventive and therapeutic interventions has been theorized from our initial observation of heterogeneous feeding preference of *S. babu babu* on both *Hemidactylus frenatus* and *Homo sapiens sapiens* from the VL-non-endemic region of West Bengal, India (Fig. 1). We questioned whether, if only the bacteria, but not the parasites, of VL-non-transmitting sand flies are inoculated into hosts, what the immune consequences would be, in the context of both pre– and post-*Leishmania* infection, as no concrete report is available on the natural transmission of VL by *S. babu babu*^63^. How will it behave, pro-parasitic or anti-parasitic? Moreover, the sand fly gut commensals may be overlapping or highly specific, depending on biotope, vector species, and host plant preferences of the vectors^64^. This work translates the epidemiological evidence of human-probing VL-non-transmitting sand fly into molecular evidence that the gut bacteria of that species, *S. babu babu,* can inhibit *Leishmania* pathogenesis in hosts.

Although the function of IL-23 has been established in autoimmune diseases^65^, it is now clear that IL-23 is important for the differentiation of IL-17-producing CD4^+^ and CD8^+^ T cells against infection^48–50,66^. Both Gram-positive and Gram-negative bacteria have been reported to induce IL-23^67^. IL-23∼IL-17 activation is required, independent of IL-12∼IFN-γ, for bacterial clearance^68^; more importantly, IL-23 is critical for long-term control against bacteria^69^. Scientific controversy remains; the rigidity of the conventional concept of IL-12∼Th1 for anti-leishmanial immunity prevails; however, recent reports strengthen the importance of independent activation of IL-23∼Th17 in protection against *Leishmania* and other protozoan diseases^48–50^. This finding wisely leveraged the heat-inactivated *H. sporothermodurans*-mediated activation of the IL-23∼IL-17 axis in favor of *Leishmania* eradication. The induction of IL-17 was greater than that of IL-12 by heat-inactivated *H. sporothermodurans*, which is justified as the exponential phase inducing IL-12, whereas the peptidoglycan after heat-inactivation inhibits both IL-12p40 and IL-12p35 expression, leading to upregulation of IL-23p19 and the generation of Th17 as reported earlier^70,71^, corroborating our observation that the heat-inactivated form is more efficient than live *H. sporothermodurans* in inhibiting drug-resistant *Leishmania*, *in vivo* (Fig. 4, Supplementary Fig.4).

IL-18, along with IL-1β, can be induced by Gram-positive bacteria by engaging the innate host immune sensors NLRP3^72^ and NLRP6^73^ inflammasomes, triggering the caspase-1 pathway, independent of IL-12, which corroborates our observation (Fig. 4). When early-acting IL-18 and late-acting IL-23 were found to be instrumental against *L. donovani*, independent of IL-12^74^, IL-1β was reported to strengthen Th17 activation in association with IL-23^51–53^. The traces of IFN-γ^+^ CD4^+^ and CD8^+^ T cells were induced by IL-18^75^ in the absence of IL-12 (Fig. 4 and Fig. 5).

VL-transmitting sand flies inoculate *Leishmania* directly into the hosts’ circulation, creating a bleeding wound with their mouthparts^76^ The success of vaccination depends on the host’s immune response, which blocks pathogens at the initial stage of entry. The host’s circulating neutrophils and monocytes are the first line of defense against invading *Leishmania*^45,54–59,77^, and specifically, monocytes act as the primary barrier against *Leishmania*, as the parasites are transferred to them by neutrophils^77^. Interestingly, the prophylactic administration of heat-inactivated *H. sporothermodurans* not only induced phagocytic capacity, but the strategy also prepared the host to resist the initial entry of *Leishmania* by inducing ROS and TNFα in circulatory CD11c^−^CD11b^+^Ly6G^−^Ly6C^hi^ monocytes (Fig. 6). The supremacy of the heat-inactivated formulation over live *H. sporothermodurans* in inhibiting parasite infection can be claimed as a novel therapeutic and/or prophylactic, with field-to-laboratory translation.

To the best of our knowledge, this is the first report of the development of a prevention and treatment against drug-resistant *L. donovani* by a bacterium, *H. sporothermodurans* [NCBI Sequence Read Archive (SRA): PRJNA1403236, whole-genome sequence], isolated from VL-non-transmitting *S. babu babu*, by the activation of the IL-23∼Th17 pathway, more importantly by impeding the establishment of *Leishmania* into the host circulation. Further clinical trials are needed to warrant (Indian patent ref: 202511041013, April 28, 2025).

## METHODS

### Ethics statement

The experimental protocols were approved by the Institutional Animal Ethics Committee (File no. IAEC/1394/2019-20/05) following ARRIVE guidelines and the Bio-safety Committee (File no. WEST16102019753/2/1.2) of West Bengal State University, Barasat, West Bengal, India. Consent was taken from the heads of the houses for indoor and outdoor collection of sand flies, from adjacent cowsheds, either by aspirators or light traps.

### Experimental animals and parasites

Male BALB/c mice are widely accepted as the experimental model for *Leishmania* research because their genetic predisposition toward Th2 immune responses makes them suitable for studying infection susceptibility that mimics human immunopathology^78^. Four to six week old male Charles-River BALB/c mice of weight (25 to 30 gm) were procured from Hylasco Biotechnology (India) Pvt. Ltd, Hyderabad, India and were reared at 25°±2°C with controlled humidity (45-50%), maintained a circadian 12-hour light/dark cycle at the animal facility of West Bengal State University, Barasat, West Bengal, India. Animals were provided with access to a standard pellet diet and water, *ad libitum*. Animal experimental protocols were pre-approved by the Institutional Animal Ethics Committee, WBSU, Barasat (File No. IAEC/1394/2019-20/05, dated 20.05.2019) in accordance with in accordance with CCSEA, Government of India and ARRIVE guidelines.

BALB/c mice were infected with antimony-resistant *L. donovani* NS2^45,79^ strain for maintenance every four weeks by serial passaging. Promastigotes were transformed from splenic aspirates of infected animals and were reared at 22-24 in a complete M199 medium with 20% heat-inactivated FCS (GIBCO) and 1% penicillin-streptomycin (Invitrogen) (details of both animal and parasite strains provided in Supplementary list 1).

### Field collection of sand flies, morphological and molecular identification

Sand flies were captured from human dwellings (mud houses) and cow sheds by using CDC Miniature Light Trap Model 512 (John W. Hock, Gainesville, FL) light traps and mouth aspirators (John W. Hock, Gainesville, FL) in different VL-non-endemic (South 24 Parganas) VL-endemic (Malda and Darjeeling) districts of West Bengal, India (Fig. 1) within the time span of October, 2019 – November, 2024 with at least 10 field visits in each districts in different seasons. Sand flies were identified morphologically and based on cibarial teeth count numbers^28,29,34^. Field-caught sand flies were euthanized by cold shock either on an ice chunk in the study field or at –20 °C for 10-20 minutes in the laboratory, with frequent monitoring to confirm they reached a stunned state. Samples were preserved in 95% research-grade ethanol for molecular biology work. Guts were immediately dissected for isolation of bacteria before long-term storage^80^.

The hexadecyltrimethylammonium bromide (CTAB) (Sigma-Aldrich) method, followed by Proteinase K (Sigma-Aldrich) treatment and phenol-chloroform-isoamyl alcohol (HIMEDIA, Sigma-Aldrich) extraction, was performed to isolate the genomic DNA from the ethanol-preserved field-caught sand flies, as we reported earlier^81^. The quantity and quality of the genomic DNA were checked using a double-beam UV-Vis spectrophotometer (Eppendorf AG, 22331 Hamburg, serial no. 6135JG004726, Germany). Molecular typing of sand flies was performed based on sequence-based analysis targeting insect-specific Cytochrome c oxidase I (COI) mitochondrial gene^82,83^.

The PCR program was as follows using primer pair: (forward primer L1440d 5’ TYTCAACWAATCATAARGATATTGG 3’, reverse primer H2123d 5’ TAWACTTCWGGRTGWCCAAARAATCA 3’) denaturation at 95°C for 5 min; followed by initial 10 cycles at 94°C (30sec), 45°C (1min), 72°C (1min), followed by 30 cycles for 94°C (1min), 51°C (1min), 72°C (1min), followed by final extension step at 72°C for 10 minutes, which was modified from the program used by Kumar *et al*. 2012 ^30,83,84^. Sanger sequencing was performed in the DNA Sequencer (ABI 3730 XL, Applied Biosystems, Waltham, MA) based on the amplified COI gene, and the retrieved nucleotide sequences were analyzed by BioEdit Sequence Alignment Editor v.7.2.5 (Supplementary list 1), followed by Basic Local Alignment Search Tool (BLAST) to compare with the available GenBank database ^81^. Multiple nucleotide sequences were aligned using the ClustalW algorithm, and genetic divergence was calculated using MEGA 11(Supplementary list 1). The Neighbour-Joining (NJ) method, alongside Kimura 2-parameter (K2P) genetic distances, was used to create a phylogenetic dendrogram and was evaluated with 1000 bootstrap replicates (details of sand flies provided in Source data file)^35,85,86^.

### Blood meal analysis

Total mRNA was extracted from each blood-fed gut of *Sergentomyia babu babu* by using the conventional TRIzol (Invitrogen) extraction method^40,45^. Synthesized cDNA was subjected to reverse transcriptase PCR amplification, based on the primer sets of *Hemidactylus frenatus*-specific COI mitochondrial gene^32,36^ designed by us and *Homo sapiens sapiens*-specific AluYB8 element DNA^37^ in the Master Cycler (Eppendorf Mastercycler Nexus Gradient, cat. 6331000017, 2020) for identification of host preferences of sand flies.

### Isolation of sand fly mid-gut bacteria and species identification

At least 30-40 surface-sterilized field-caught live female sand flies were taken after euthanasia, followed by three washes with 1X Phosphate Buffered Saline (PBS). The midgut was dissected following a sterile protocol, homogenized in 1X PBS, and 40 µl of the homogenate was plated on a BHI agar (Sigma-Aldrich) plate from a pool of 10 sand flies. The plates were incubated in a 36±1°C incubator for 24-48 hours^22,38^. Distinct colonies were picked for inoculation into liquid BHI broth (Sigma-Aldrich), incubated overnight on a rotary shaker (250 rpm, 36±1°C) for multiplication, and replicas were taken for either DNA isolation or preservation at 70°C^87^. Sanger sequencing of the amplified bacterial housekeeping 16S rRNA gene, followed by sequence analysis, was performed as described earlier. Specimens with ≥97% nucleotide sequence similarity to their closest relatives were considered the same species (details of bacteria provided in Source data file).^81,87^

### Infection, therapeutic and prophylactic schedule, and parasite load determination

*H. sporothermodurans* was identified by whole-genome sequencing (WGS) of the bacteria and was grown in BHI broth overnight at 36±1°C for 24 hours. The 5 × 10^6^ CFU of *H. sporothermodurans* was determined using the standard curve at an optical density (OD) of 600nm (OD_600nm_), as per our laboratory-developed protocol. For the heat-inactivated bacterial formulation, 5 × 10^6^ CFU of *H. sporothermodurans* was resuspended in 1X PBS and incubated at 70°C for 40 minutes^88^. Heat-inactivated *H. sporothermodurans* (5 × 10^6^ CFU / animal) in 50µl 1X PBS formulation was administered, sc, on the 1st, 3rd, 5th, 7th, and 14th day, either before (prophylactic) or after (therapeutic) the infection of male BALB/c mice (5 animals per group) with drug-resistant *L. donovani* NS2 ^45,79^ intravenously^40,45^. For the appropriate dose, route of administration, and comparative efficacy, the variable doses (6×10^6^ CFU, 5×10^6^ CFU, 4×10^6^ CFU, and 3×10^6^ CFU/ animal), variable routes (subcutaneous, intramuscular, and intradermal) in both live and heat-inactivated formulations, and variable adjuvants (*Bacillus* Calmette-Guérin and *Mycobacterium indicus pranii*) were chosen. Control groups received no treatment. Experimental animals were euthanized either at the end of the study or upon reaching humane endpoints in accordance with American Veterinary Medical Association guidelines. Mice were anesthetized with a 2-3% isoflurane/O2 mixture to collect whole-blood samples or were exsanguinated via cardiac puncture. Following exsanguination, confirmation of death occurred via cervical dislocation. Terminal tissues were collected rapidly for further processing according to the needs of the experiments^89^. Bone marrow, spleen, and liver samples were collected one month post-infection, either preceded (prophylactic) or followed (therapeutic) by a 15-day treatment schedule to estimate parasite burden. Parasite burden was determined from Giemsa (Sigma-Aldrich)-stained splenic and hepatic tissue micrographs applying Stauber’s formula^40,45^. More precisely, *L. donovani*-specific 18S rRNA was quantified to determine infection severity in both bone marrow and spleen, followed by normalization to murine GAPDH in Real-time PCR, as previously described ^40,45^.

### Immune activation and inhibition in initial parasite entry and establishment

#### Real-time PCR for cytokine induction

Splenocytes were collected from each group (naïve, infected, and treated) of experimental animals, cultured for 6 hours, stimulated by 25 *µ*g/ ml of soluble crude *Leishmania* antigen at 37 C and in a 5% CO_2_ environment. The mRNA was extracted as per the conventional TRIzol method^40,45^, followed by the cDNA preparation and estimation of induction of cytokines (IL-17, IL-1β, IL-12p40, IL-12p35, IL-6, IL-23, IL-18, IFN-γ, IL-10 and TGF-β) in respect to normalized murine GAPDH as per our regular protocol (details provided in Supplementary table 20)^40,45^, using specific oligonucleotide primers and iTaq™ Universal SYBR Green Supermix (Biorad) in CFX opus@96.

#### Expansion of IFN-γ^+^ and IL-17^+^ splenic T cells

Splenocytes were collected from one-month post-infection animals, followed by a therapeutic schedule, and cultured overnight in the presence of 25 µg/ml crude *Leishmania* antigen at 37 C and 5% CO2 after RBC lysis. Cells were harvested and stained for flow cytometry analysis as per the standard protocol of our laboratory ^40,45,90^. In brief, the dead cells were excluded by using live/dead dye for 15 minutes at (RT) and purified rat anti-mouse CD16/CD32 (Fc blocker, catalogue number 553142, BD) was used to block the cells for non-specific binding, followed by their staining with fluorochrome-conjugated antibodies (details provided in Supplementary list 2) with respect to fluorescence minus one (FMO) controls and the antibody specific isotypes (Supplementary Fig. 5 and 6) for induction of IFN-γ and IL-17, both on CD4^+^ and CD8^+^ T cells^40^ in BD FACSVerse^TM^, and data matrices were evaluated by BD FACSuite software.

#### Phagocytic uptake during entry of Leishmania in host circulation by Ly6G^−^Ly6C^hi^ monocytes

Experimental animals were immunized with heat-inactivated *H. sporothermodurans* as per the prophylactic schedule (5×10^6^ CFU per animal). Promastigotes of drug-resistant *L. donovani* NS2^45,79^ were stained with VPD450 (violet proliferation dye450, cat# 562158, BD Biosciences) as we reported earlier^45^ and released (2×10^7^ VPD450^+^ *Leishmania*) into the circulation of the immunized mice, iv, 3 hours after the last dose of immunization^45^. Naïve, only immunized (no infection), and immunized (infection) groups were used as controls. RBC-lysed white cell packs were used for fluorochrome staining and flow cytometry analysis as per the regular protocol of our laboratory ^40,45^. VPD450^+^*Leishmania* was quantified in the specific cell population of circulatory CD11c^−^CD11b^+^Ly6G^−^Ly6C^hi^ monocytes (Supplementary list 3) to estimate phagocytic activity in different experimental groups at 3 hours post-infection.

#### Induction of TNF-α and ROS in circulatory Ly6G^−^Ly6C^hi^ monocytes

Cells were stained with DHR123 (Sigma-Aldrich) and anti-mouse TNF-α as per the regular protocol of our laboratory^40,45^. ROS (DHR123^+^) and TNF-α were measured in the specific cell population of circulatory CD11c^−^CD11b^+^Ly6G^−^Ly6C^hi^ monocytes to assess leishmanicidal activity (Supplementary list 3).

## Statistical analyses

The statistical measure (mean±SEM) of minimum of 5 biological replicates combined from two independent experiments was used to interpret all experimental data, followed by an unpaired two-tailed Student’s t-test or one-way analysis of variance (ANOVA) to calculate the p-value. Graphical representations were generated using GraphPad Prism software (version 8.0). A p-value of <0.05 was considered statistically significant.^40,45^

## Data availability

All the nucleotide sequences were retrieved from Sanger sequencing and whole genome sequencing data were deposited in NCBI GenBank and NCBI Sequence Read Archive (SRA): Accession numbers [PV362839, PV366908, PV362840, PV387274, PX830396, PV650449, PV363009, PV363008, PV362971, PV362972, PV363010, PX830395, PV650444, PV367238, PV412603, PV650854, PV362973, PV650450, PV650853, PV650785, PV650299 and PRJNA1403236]. All datasets, links to NCBI accession numbers, and individual data are provided in the MS, supplemental information file, and source data file, and are publicly available (doi: 10.17632/hbps3wnzdd.1). Additional information on the other data analyses is available upon request from the lead contact. The preprint version of the MS has been published (……).

## Code availability

This article did not report any original code.

## ACKNOWLEDGMENTS

We thank the Vice Chancellor of West Bengal State University, Barasat, for providing the research infrastructure for this work. We acknowledge DST-FIST, Govt. of India (Ref. SR/FST/LS1-001/2014), DBT-BOOST, Govt. of West Bengal (Ref. 49[11]/BT [Estt]/1P-4/2013 [Part-1]), and the Department of Higher Education Govt. of West Bengal (Ref. 1282 (Sanc)/HED-12011(48)/22022-UNIVSEC-Dept. of HE) for providing funds for the flow cytometry and real-time PCR facility in the Department of Zoology, WBSU, Barasat. SK received the fellowship from UGC, Govt. of India (Ref no. 745/(CSIR-UGC NET DEC.2017); PM received the fellowship from UGC, Govt. of India (Ref no. NTA 201610092746); SN received the fellowship from Indian Council of Medical Research Govt. of India (Ref no. Fellowship/100/2022-ECD-II); SM received the fellowship from UGC, Govt. of India (Ref no. 211610055151); and KR received the fellowship from DST-INSPIRE, Govt. of India (Ref no. DST/INSPIRE/03/2016/000117).

## FUNDING

This work was supported by the Indian Council of Medical Research, Government of India (Ref: 6/9-7(214)/2019-ECD-II, dated 01.10.2019).

## AUTHOR CONTRIBUTIONS

**Suman Karmakar:** Visualization, Methodology, Investigation, Formal analysis, Data curation, Writing-first draft; **Pritam Mandal**: Methodology, Investigation, Formal analysis,; **Supriya Nath**: Methodology, Investigation, Formal analysis; **Soumyadip Mukherjee**: Methodology, Investigation, Formal analysis; **Kamalika Roy**: Methodology, Investigation, Formal analysis; **Mintu Karan**: Investigation; **Sharmistha Paul**: Investigation; **Sondipon Chakraborty**: Investigation; **Ashif Ali Sardar**: Resources (assisted sand fly sampling, Malda district), **Mithun Sarkar**: Resources (assisted sand fly sampling, Darjeeling district), **Pradyumna Patra**: Investigation, Resources (assisted sand fly sampling, South 24 Parganas district); **Chiranjib Pal:** Conceptualization, Visualization, Writing-review & editing, Supervision, Funding acquisition, Project administration.

## DECLARATION OF INTERESTS

The authors declare no known competing financial interests or personal relationships that could have influenced the work reported in this paper.

## MATERIALS AND CORRESPONDENCE

All information related to resources and reagents will be available from the corresponding lead contact, Chiranjib Pal.

## DECLARATION OF GENERATIVE AI AND AI-ASSISTED TECHNOLOGIES

The authors did not use any AI technologies to prepare the manuscript. Both AI-generated-only and AI-generated text that was AI-paraphrased were detected at 0% by the University licensed plagiarism software (Turnitin and DrillBit-Extreme through INFLIBNET, UGC, Govt of India).

## PATENT STATEMENT

An Indian patent has been filed jointly by the Indian Council of Medical Research, Government of India and West Bengal State University, Barasat, West Bengal, India, entitled ‘Formulation for the Treatment and Prevention of Leishmaniasis’ [Ref: 202511041013, April 28, 2025].

## SUPPLEMENTAL INFORMATION

All supplementary figures, tables, and reagents, including original gel images, have been provided in a separate file.

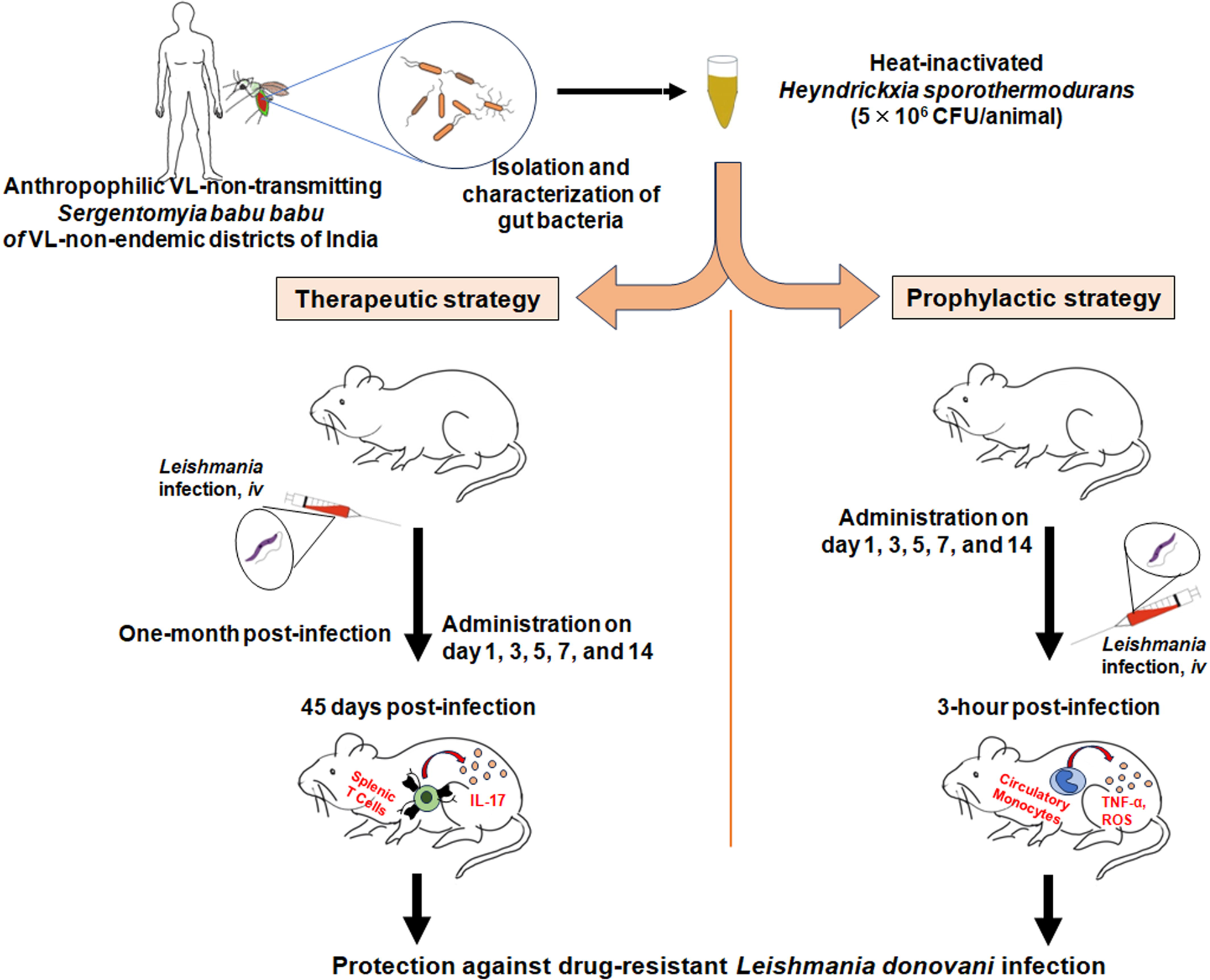

## Notes

https://10.17632/pxc6jbw7t3.1

### Competing Interest Statement

The authors have declared no competing interest.

